# Appetitive learning relies on octopamine and dopamine in ants

**DOI:** 10.1101/2021.04.19.438615

**Authors:** Maarten Wissink, Volker Nehring

## Abstract

Associative learning relies on the detection of coincidence between a stimulus and a reward or punishment. In the insect brain, this process is thought to be carried out in the mushroom bodies under control of octopaminergic and dopaminergic neurons. It was assumed that appetitive learning is governed by octopaminergic neurons, while dopamine is required for aversive learning. This view has been recently challenged: Both neurotransmitters seem to be involved in both types of memory in bees and flies. Here, we test which neurotransmitters are required for appetitive learning in ants. We trained *Lasius niger* ant workers to discriminate two mixtures of linear hydrocarbons and associate one of them with a sucrose reward. We analysed the behaviour of the trained ants using machine learning and found that they preferred the rewarded odour over the other, a preference that was stable for at least 24 hours. We then treated the ants before learning with either epinastine, an octopamine receptor blocker, or with flupentixol, a dopamine receptor blocker. Ants with blocked octopamine receptors did not remember the rewarded odour. Octopamine signalling is thus necessary for the formation of appetitive memory. In contrast, ants with blocked dopamine receptors initially learned the rewarded odour but failed to retrieve this memory 24 hours later. Dopamine is thus required for long-term memory consolidation during appetitive conditioning, independent of short-term memory formation. Our results show that appetitive learning depends on both octopamine and dopamine signalling in ants.

## Introduction

There is nothing stable in the world; those that can adapt to changes will prevail. When the profitability of food sources changes over time, it would be advantageous to quickly learn which ones are currently most rewarding. Individuals can do so by associating the food with other stimuli such as odours or colours, i.e. they establish a predictive relationship between two independent cues from the environment (Giurfa, 2007). When a neutral stimulus, like an odour, is paired with a biologically relevant stimulus (food, the unconditioned stimulus – US), the animal may in future react to the previously neutral odour (the now conditioned stimulus – CS), as if it were the US. In the case of a bee searching for food, CS and US are processed by two different pathways. The CS-pathway starts at the odorant receptors (ORs) in the antenna. Output from the OR neurons is first processed in the antennal lobe (Giurfa, 2007). From there, projection neurons carry the information to the mushroom bodies, where learning occurs. The US-pathway begins at gustatory receptors on the mouth parts. Neurons then project to the subesophageal ganglion, from where the octopaminergic VUMx1-neuron projects to the antennal lobe and the mushroom bodies (Giurfa, 2007; Galizia and Sachse, 2010; Perry and Barron, 2013). CS and US converge in the Kenyon Cells of the mushroom body, which act as coincidence detectors. In their normal state, they would not relay a specific odour to the higher brain centres through mushroom body output neurons. However, after an odour coincided with an US, the Kenyon cells will relay the odour as a now conditioned stimulus, even in absence of the US.The reliance of the US signalling on the octopaminergic VUMx neuron suggests that without octopamine (OA) signalling, the US cannot reach the MB and thus insects cannot learn. Indeed, blocking OA receptors in crickets, flies, and honeybees prevented appetitive memory formation and memory retrieval (Schwaerzel et al., 2003; Galizia and Sachse, 2010), while OA injections could replace the US (Hammer and Menzel, 1998). Similar experiments blocking dopamine (DA) receptors had no effect on appetitive learning but prevented individuals from associating a punishment, such as an electric shock, with odours or colours (aversive conditioning; Schwaerzel et al., 2003; Vergoz et al., 2007; Mizunami and Matsumoto, 2017). It was thus assumed that aversive and appetitive memory are formed in different modules of the antennal lobes that use different neurotransmitters (Hige et al., 2015). To this day, this seems to hold true for crickets (Mizunami and Matsumoto, 2017), but newer data suggest that aversive conditioning relies on both OA and DA signalling in honeybees and fruit flies (Agarwal et al., 2011; Claßen and Scholz, 2018; Manicini et al., 2018). The use of *Drosophila* GAL4 lines that can be used to knock out specific neurons in the MB has allowed to paint a much more detailed picture of learning and memory retrieval: In flies, learning in the MB appears to be organized in modules that each consist of Kenyon cells, mushroom body output neurons, and dopaminergic neurons that modulate the valence of the modules (Berry et al., 2012; Hige et al., 2015.) Some of the modules organize attraction to odours, and others repulsion (Rohwedder et al., 2016). The modules further differ in how long the memory lasts – from a few seconds to several hours (also in honeybees: Menzel, 2014). The control of both appetitive and aversive learning by dopaminergic neurons also means that without dopamine signalling, both types of learning should be hampered, which matches the behavioural evidence from pharmacological receptor blocking and knockouts in *Drosophila* and honeybees (Berry et al., 2012; Perisse et al., 2013; Perry and Barron, 2013; Mancini et al., 2018; Sabandal et al., 2020).

One way to hinder dopamine- and octopamine-related signalling is to block their respective receptors pharmacologically. In insects, epinastine is a specific OA-receptor blocker which abolishes the OA related cAMP production in the mushroom bodies (Roeder et al., 1998). Epinastine seems to specifically target the AmOA1 receptor in honey bees (Beggs et al., 2011), the OA1 receptor in crickets (Awata et al., 2015), the DmOA3 receptor in *Drosophila* (Qi et al., 2017), and the and NcOA2B2 receptor in the green rice leafhopper *Nephotettix cincticeps* (Xu et al., 2020). In contrast, flupentixol largely targets dopamine receptors, such as the AmDOP2 receptor in the mushroom bodies of honey bees (Mustard et al., 2003; 2010; Beggs et al., 2011).

Just like honey bees, ants are social insects for whom learning is important in many contexts (Bos et al., 2010). Ants can learn odours of rewarding food sources and also have to learn the specific odour of their own colony (Neupert et al., 2018). Indeed, ants can perform similar learning tasks as bees and other insects (Bos et al., 2010, Guerrieri and d’Ettorre, 2010; Fernandes et al., 2018; Piqueret et al., 2019). In addition to floral odours that might signal food sources, ants were successfully conditioned to hydrocarbons that play an important role in social interactions (Bos et al., 2012, Sharma et al., 2015). For example, hydrocarbons can serve as alarm and queen pheromones, and are used to discriminate nestmates from non-nestmates (Leonhard et al., 2016). While it has been found that protein synthesis is important for the formation of the long term memory in *Camponotus* ants (Guerrieri et al., 2011), we know nothing about the neural circuits of learning in the ants, nor of the role different neurotransmitters play in ant learning.

Here, we study reward learning the black garden ant *Lasius niger*. First, we established an assay that trained the ants to associate a mixture of linear hydrocarbons to a sugar reward. We tested the memory retrieval after 5 minutes and after one day using a deep learning algorithm on videos of reward-searching ants. In a second step, we administered epinastine or flupentixol via either the food or topical application, to block OA and DA receptors and to test if a breakdown of the respective neural pathways would prevent the ants from forming short- and long-term memories of the odour-reward association.

## Material and Methods

We trained ants to discriminate two mixtures of n-alkanes, one of which was presented with a sugar reward and the other was unrewarded, during six rounds of training trials. After that, we ran retention tests without sugar reward to see whether the ants now preferred the previously rewarded odour (conditioned stimulus - CS+) over the unrewarded odour (CS0). A first retention test was conducted ten minutes after the final learning trial to test the short term memory capabilities. The second retention test was performed one day later to test the (early) long term memory (Menzel, 1999).

Before the learning trials, we submitted the ants to treatments with blockers of octopamine receptors, dopamine receptors, or controls. The drugs were administered either with food (set 1) or by topical application on the mesothorax (set 2). The two experimental sets also differed in the way we recorded the data (manual vs. automated recording, see Experimental Setup).

### Experimental Organism

Colonies of black garden ants *Lasius niger* (Linnaeus, 1758) were collected in the years 2017 and 2019 in Freiburg, Germany. They were transported to the laboratory and kept in plastic boxes at 20°C, a 12:12 light:dark cycle, and were fed with meal-worms (*Tenebrio molitor*) and honey. Test tubes filled with water, cotton wool, and an aluminium foil wrapping were placed in the boxes as a refuge and a water source. The boxes’ walls were lined with Fluon® (AGC Chemicals Europe, Ltd.). Around two weeks before an experiment, the ants were deprived of honey and only fed mealworms, in order to motivate them to forage for carbohydrates. We used four different source colonies for the experiments in set 1, and 11 different source colonies for set 2.

### Stimuli

Two mixtures of three long-chain linear alkanes each were used as the two conditioned stimuli (CS). The first mixture (CS-mixture A) was n-octadecane (n-C18), n-heneicosane (n-C21), and n-heptacosane (n-C27). The second (CS-mixture B) was n-eicosane (n-C20), n-docosane (n-C22), and n-pentacosane (n-C25; all Sigma–Aldrich, Steinheim, Germany). The hydrocarbons were solved in n-pentane (10 μg/ml) that would evaporate before the experiments. In a preliminary experiment, we used the retention test setup (see below) with untrained ants to see if they preferred any of the two odour mixtures over the other. The untrained ants spent similar times with the two odour mixtures (n = 30, wilcoxon test V = 253, p = 0.69; Figure S1). A 0.5 μL drop of sugar water (50% w/w) or pure water were used as a positive unconditioned stimulus (US+) and neutral unconditioned stimulus, respectively.

### Experimental Setup

For the first set of experiments (feeding experiment) the test arena was the top of a petri dish (90 mm in diameter). Another petri dish lid was used as a roof, and a hole (40 mm in diameter) was cut into the roof to allow placing the ants into the arena. The ants were recorded from a 45° angle by a Nikon D90 with a 35mm lens, which was placed in front of the arena on a tripod. For the second set of experiments (topical application), an acrylic glass cylinder (100 mm diameter – 100 mm height) was used as an arena. The arena was covered by a cardboard box lined with aluminium foil to exclude external stimuli and to allow for even illumination by two light sources above the arena. The ants were videotaped with a SM-P600 tablet computer at 1.5x magnification, the lens of which was placed directly above the centre of the arena.

For both sets of experiments, the walls of the arena were coated with Fluon to prevent the ants from climbing. The floor was lined with filter paper that was evenly divided into quarters. A microscope cover slip (24×24 mm) was placed in the centre of each quadrant (Fig. 1A). In each learning and test trial, one of the cover slips was coated with 20 μL of odour A, another with odour B, and the remaining two with pure pentane. The solutions were always applied to the edges of the cover slips. After every trial, the filter paper and cover slips were replaced with new ones.

**Figure 1.**
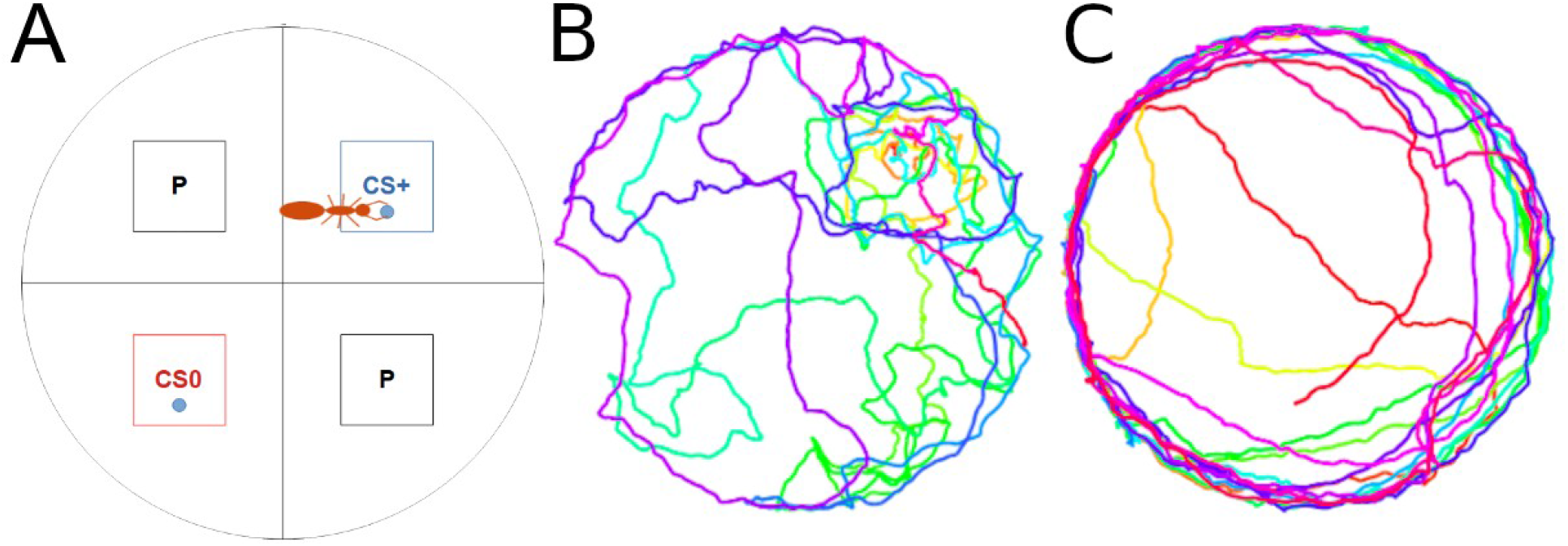
A) Design of the experimental arena used for the learning experiments. The arena was divided into four quadrants, each with a cover slip in it. One of the cover slips was coated with odour A, another one with odour B. During the learning experiments, one of the odours was paired with sucrose solution and the other with water. B, C) Traces of ants during the retention tests with the rewarded odour in the top right quadrant. The ant in B inspected the CS+, the ant in panel C did not show any particular interest in the previously rewarded odour. Colours show how the ants moved from the beginning of the test (red) to the end (blue).

### Learning Trials

The learning procedure was adapted from Bos et al.’s (2010, 2012) work on ant learning. From the main colony, around 30 ants that were moving outside of the nest tubes were separated into a box equipped with nest tubes like the source colonies were. Here, ants could perform trophallaxis between the trials, to reduce their sugar storage and increase the motivation to forage again. The ants were marked on the gaster with a small dot of paint (Edding No. 751), so that it was possible to individually track them between the learning trials.

For each learning trial, 0.5 μl of sugar solution was added to the cover slips with one of the odours (CS+), while water was added to the slip with the CS0. For each ant, the same odour was rewarded in each learning trial, but which odour was rewarded differed randomly between ants. An individual ant was then lifted gently into the arena and the time was measured until it began drinking the sugar solution. Then, the ant was removed to prevent it from consuming too much sugar solution, and placed back into the experimental colony where it could perform trophallaxis with the nestmates. After six minutes, when trophallaxis was always complete, the ant was brought back into the arena for the next learning trial. In total, there were six learning trials per individual.

If an ant had not found the sugar solution within 3 minutes in set 1, she was removed from the learning arena and brought back into the experimental colony until the next learning trial. Ants that could not find the sugar solution in two or more learning trials were excluded from the analysis. The process was similar for set two, but we gave the ants only two minutes for the second through sixth learning trial, because the results from set 1 showed that times longer than 2 minutes are very rare in the later trials. We began set 1 with 180 ants across the six treatments and excluded 14 of those. In set 2, we had to exclude 9 out of 189 ants. The highest proportion of excluded ants came from flupentixol treatments in both sets (Table S1). There was no difference between treatments when we fed the receptor blockers (χ^2^_5_ = 8.5, p = 0.13, p-value calculated by Monte Carlo simulation). In the topical application experiment, the treatments differed in the likelihood of ants failing (χ^2^_3_ = 10.7, p = 0.02), although overall only 2 ants were excluded, both from the flupentixol treatment.

We tested whether the ants became quicker to find the sugar solution during the consecutive learning trials with a poisson MCMC glmm, with time until finding the sugar as the dependent variable and trial number as a continuous predictor, for each treatment separately. We entered the colony ID and individual ant ID as random factors into the model using the R package MCMCglmm with 50000 MCMC iterations and otherwise default settings (R version 4.0.3 – R Core Team, 2020; Hadfield, 2010). In set 2, we observed the ants for only 2 minutes in learning trials 2-6, as opposed to 3 minutes in trial 1. To exclude that the few ants taking longer than 2 minutes in trial 1 already cause a downward trend in the learning times across trials, we set the times of all ants that took longer than 2 minutes to 2 minutes for the glmm but left all times shorter than two minutes as they were.

### Retention Tests

When the ants had finished trophallaxis after the sixth learning trial, a first retention test was conducted. Retention tests resembled training trials but no sugar solution or water was added to any of the cover slips. The cover slips with the CS+ and CS0 odours were in opposite quadrants (Fig. 1A). Each ant was recorded for two minutes. Then, a drop of sugar water was placed on the cover slip with the learned odour to prevent extinction before the retention trials on the following day. When the ant had finished to eat, it was lifted back into the test colony. The position of the cover slips was changed and the ant was recorded a second time. The duration spent in the different quadrants was averaged over the two tests for each ant. On the next day, the same ants were tested again to see if they could still remember the CS+. The marked ants were kept in isolation over night together with the other ants from the same colony that were treated on this day. The interval between the two test trials was 15-23 hours.

The time the ants spent in the four quadrants was measured from the video recordings. In set 1, times were measured manually using the program etholog (Ottoni, 2000). For the second set, the deep learning algorithm DeepLabCut (Mathis et al. 2018) was used. The algorithm was trained to spot the ant and record its coordinates using 200 frames from each of 20 pre-recorded videos, in which the position of the ant was marked manually. Then, the algorithm went over these frames 200,000 times to perfect its ant recognition performance. After this, the first videos from the experiments were processed with the algorithm and manually checked for errors. Only few errors were detected and then used to further improve the algorithm, which was virtually perfect after ca. 10 videos. In total, 756 videos were used for data analyses.

From the ant’s coordinates, we could calculate in how many frames the ant was identified in each of the four quadrants, and thus how much time it spent there. We directly compared whether the ants spent more time on one of the two odour mixtures using paired wilcoxon tests. In addition, we calculated and visualized (ggplot2 – Wickham, 2016) a preference index as an intuitive measurement of preference and hence learning performance based on the times (t) on the different odours (Bos et al. 2010, 2012):

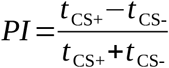

The index ranges from 1 (preference for the CS+) to −1 (preference for the CS0). If the ant spent an equal amount of time on both odours, the PI would be 0.

### Pharmalogical Manipulations

To test in how far neurotransmitters are involved in appetitive learning, the ants were treated with one of two different receptor blockers before the learning trials: epinastine, an octopamine receptor blocker (Kamhi et al., 2015), or flupentixol, a dopamine receptor blocker (Agarwal et al., 2011).

For the first set of experiments, ants were fed honey infused with the blocker (20 mM of epinastine hydrochloride or 50 mM of flupentixol hydrochloride) either 1-3 hours, 5-8 hours, or 17 hours prior to the experiment. The ants were individually fed with a 1 μL drop of this solution. The different intervals were chosen because it was not clear how long it takes for the drugs after feeding to become effective. The honey with the drug was coloured with neutral red to control if the ants had indeed ingested the honey. Only ants with a red staining in their gaster were chosen for the experiments. When the ants had finished eating, they were kept in groups with other ants that had received the same drug, until the start of the learning trials. Immediately before the beginning of its learning trial, each ant was marked with the paint marker and moved back to a subcolony of ca. 30 ants set up for this purpose. Control ants were fed pure honey instead of blocker-infused honey.

For the second set of experiments, the drugs were dissolved in dimethylformamide (DMF), and a 0.5μl droplet directly applied to the ant’s cuticula on the mesothorax (Barron et al., 2007). The ants were allowed to recover from the treatment for 15 minutes in isolation, after which they were paint-marked and returned 5 minutes later to the subcolonies as described above. Another 10 minutes later, the first learning trial began. For epinastine, the two dosages were 20mM or 100 mM in DMF, corresponding to doses of 1.8ng/μg and 9.1ng/μg of ant body weight. For flupentixol, the dosage was 250 mM (39.3ng/μg). Control ants received pure DMF.

We compared the preference indices (PI) among treatments with MCMC glmms with treatment as the only predictor, so that we could test for each treatment whether it differed from the control. Colony ID was entered into the model as a random factor. In the same manner, we compared the walking speed of ants, calculated from the coordinates generated during set 2, among the different treatments.

## Results

### Set 1: Feeding of receptor blockers

Over the successive learning trials, control-treated ants became quicker in finding the sugar reward (Fig. 2, poisson glmm p = 0.04, n = 30 ants), indicating that they learned to associate the rewarded odour with the sugar reward. This was also true for all treatments with receptor blockers (epinastine 1-3h p < 0.001, n = 27 ants; epinastine 5-8h p < 0.001, n = 29 ants; epinastine 17-26h p = 0.02, n = 25 ants; flupentixol 1-3h p = 0.04, n = 26 ants, Fig. 2), with the exception of the ants that were treated with flupentixol 5-8h before the first learning trial (p = 0.50, n = 21 ants).

**Figure 2:**
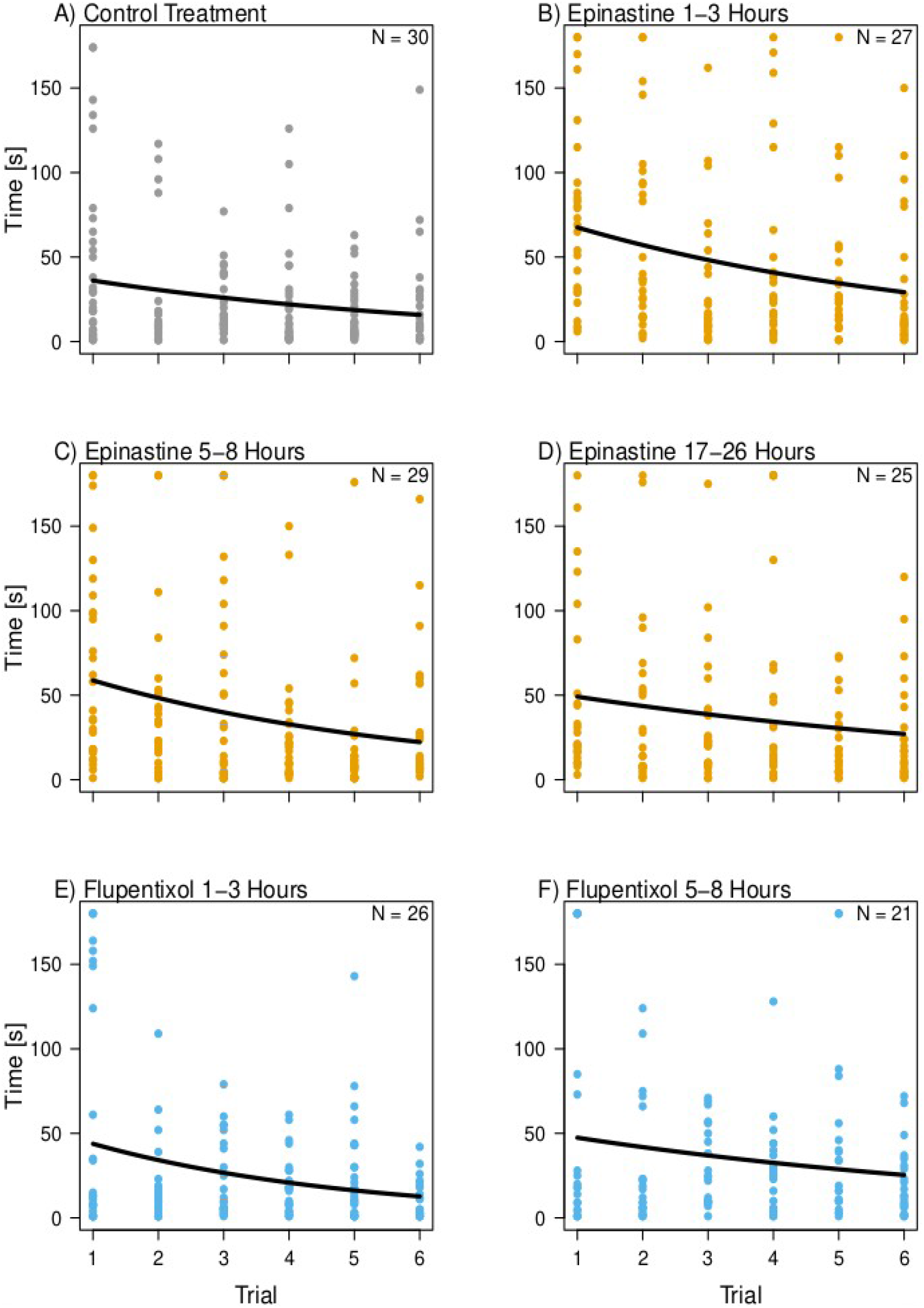
The time it took ants to find the sugar reward in six consecutive learning trials. Ants were either control treated (a), or received epinastine (b-d) or flupentixol (e, f) with their food at different intervals before the first learning trials (1-3 hours, 5-8 hours, 17-26 hours). Individual data points are plotted; the lines are poisson regressions.

In the first retention test 10 minutes after the last learning trial, the ants were able to differentiate between CS+ and CS0, independent of treatment (Figure 3A, wilcoxon-tests p < 0.05). Only the ants that were fed with flupentixol 5-8h before the treatment had difficulties to differentiate between CS+ and CS0 (wilcoxon test p = 0.052). In most treatments, the preference indices were indistinguishable from that of control ants (glmms p > 0.16; Tab. S2 for details), but the PIs of ants treated with flupentixol 1-3 hours before the learning trials were lower than those of control ants (glmm p = 0.045), and there was a similar trend for the preference indices of ants treated with epinastine 5-8 hours before the learning trials (p = 0.07). On the second day (Figure 3B), the control ants and some of the epinastine-treated (5-8 hours and 17-26 hours before the learning trials) ants still differentiated between CS+ and CS0 (wilcoxon test p < 0.05). In contrast, ants that were treated with epinastine 1-3 hours before the experiment, and the ants from both flupentixol treatments were not able to differentiate between CS+ and CS-(wilcoxon tests p > 0.05). However, there was no difference in the preference indices between the treatments (glmm p > 0.5; Tab. S2), with the exception of ants treated with epinastine 1-3 hours before the learning trials, for which there was a trend of the PIs being lower than those of control ants (p = 0.06).

**Figure 3:**
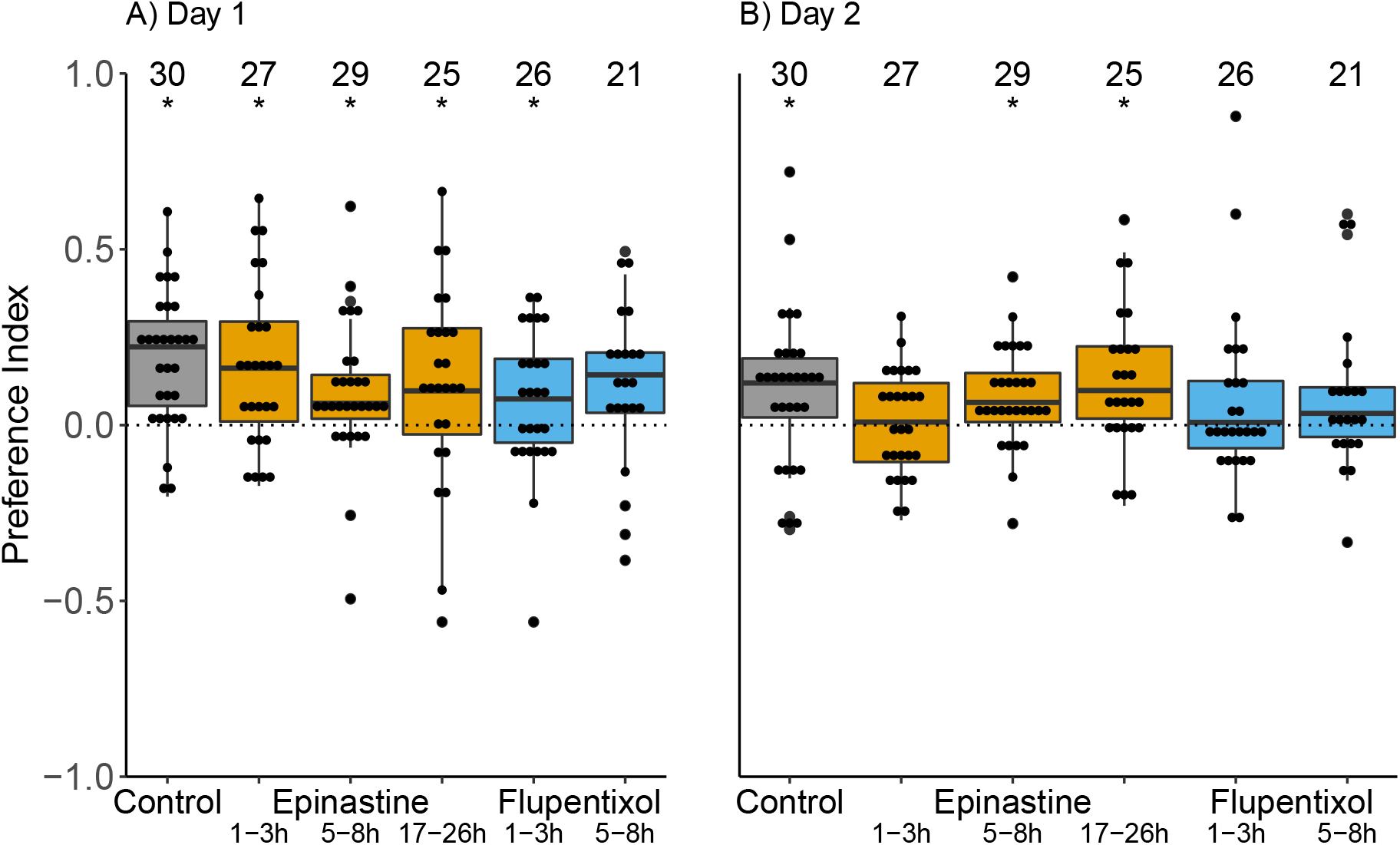
Preference for the previously rewarded odour (CS+) during retention tests with ants that received control treatment, epinastine, or flupentixol through the food. Each ant completed two retention tests without reward, one five minutes after the sixth learning trial (A), and a second test one day later (B). The ants could move in an arena in which one quadrant was marked with the CS+ odour and another one with CS0. The dotted lines indicate the null hypothesis of ants spending equal time on the CS+ and the CS0; positive values indicate that the ants spend more time in the quadrant marked with the CS+. Numbers above the boxes are sample sizes, * means that the times spent on CS+ and CS0 differed (wilcoxon test p < 0.05). For each treatment, standard boxplots and indices of individual ants are plotted.

### Set 2: Topical application

The exact dosage of the receptor blockers is difficult to control when feeding the drugs, because the ants may sometimes ingest more and sometimes less of the food. We thus performed a second set of experiments where we topically applied precise doses of the blockers 30 minutes before the learning trials. During the learning trials, the time it took the ants to find the sugar solution decreased for the control-treated ants (poisson glmm p = 0.010, n = 31 ants; Fig. 4A). This effect was also evident for ants that had been treated with 100mM epinastine (p < 0.001, n = 10 ants) or flupentixol (p = 0.001, n = 30 ants) previous to the learning trials (Fig. 4B,C). Note that we could only analyse ten of the epinastine learning trials with high dose and none of the ones with low dose.

**Figure 4:**
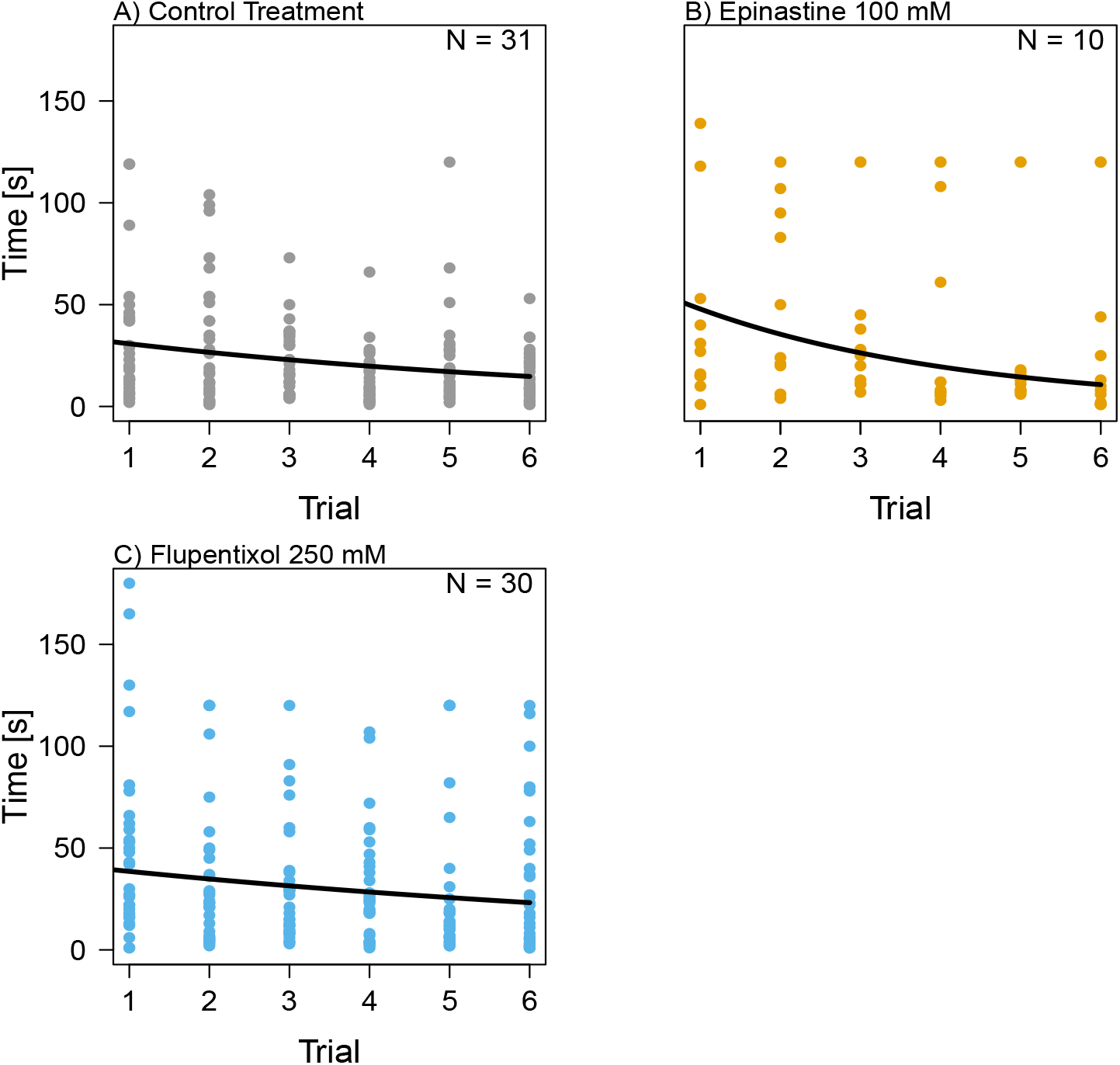
The time it took ants to find the sugar reward in six consecutive learning trials. Ants were either control treated (a), or topically treated with 100mM epinastine (b) or 250mM Flupentixol (c). Lines are from poisson regressions.

In the first retention test, the ants that had been treated only with DMF control preferred the CS+ over the CS0 (Fig. 5A; wilcoxon test p < 0.001). The same was true for ants that had been treated with 20 mM epinastine (wilcoxon test p < 0.001) or flupentixol (wilcoxon test p < 0.001). The ants that had been treated with 100 mM epinastine did not differentiate between the CS+ and CS0 on the first day (wilcoxon test p = 0.83). The preference indices of the 100mM epinastine treatment were lower than those of the control ants (glmm p < 0.001, Tab, S3), but those of the 20mM epinastine treatment (p = 0.45) or the flupentixol treatment (p = 0.18) were not.

**Figure 5:**
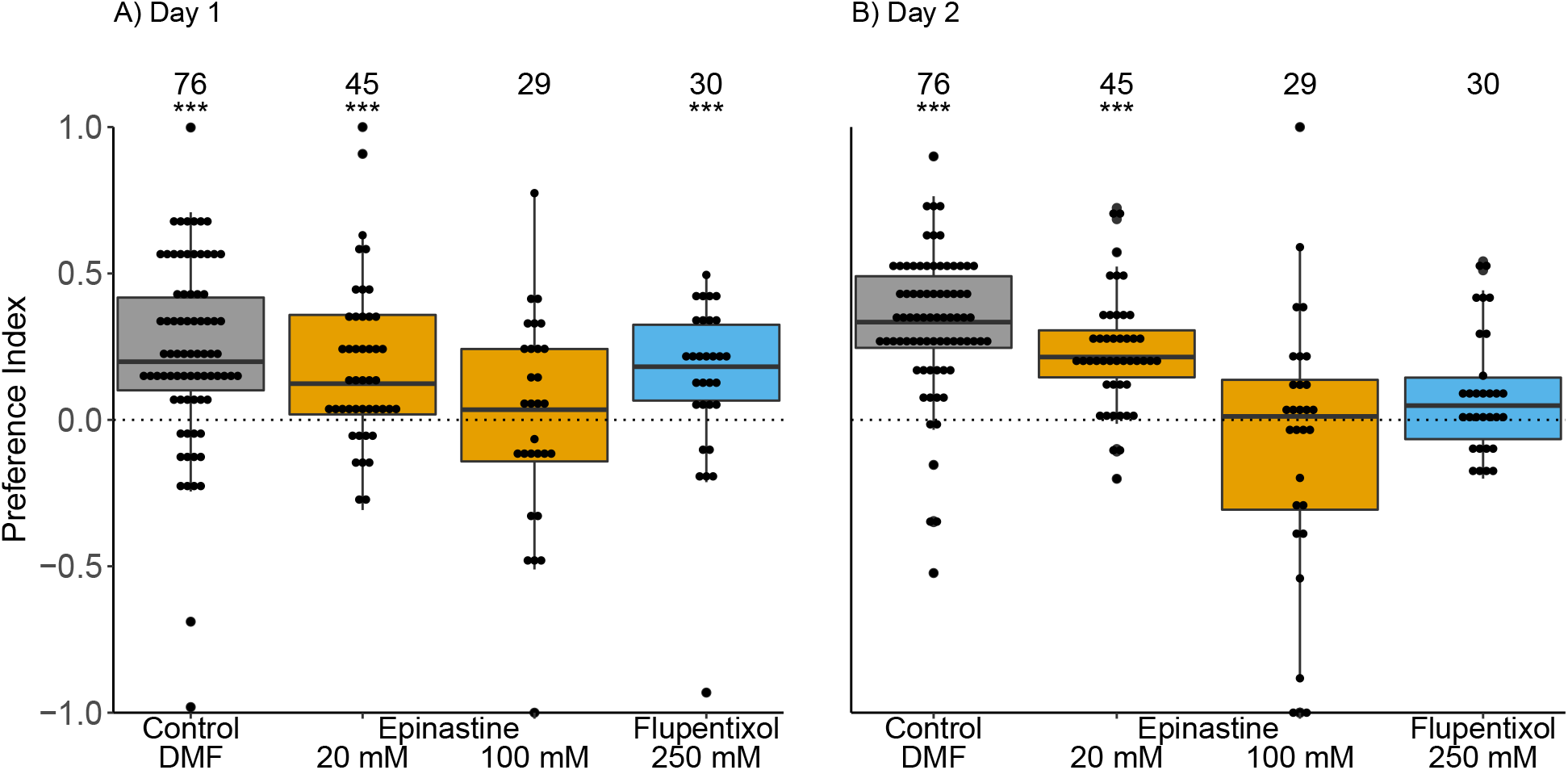
Preference for the CS+ during retention tests with ants that received control treatment, epinastine, or flupentixol through topical application. Each ant completed two retention tests without reward, one five minutes after the sixth learning trial (A) and a second test one day later (B). The ants could move in an arena in which one quadrant was marked with the CS+ odour and another one with CS0. The dotted lines indicate the null hypothesis of ants spending equal time on the CS+ and the CS0; positive numbers indicate that the ants spend more time in the quadrant marked with the CS+. Numbers above the boxes are sample sizes, stars indicate significant differences between the time spent on CS+ and CS0 (wilcoxon test; *** p <0.001).

One day later, the preference for CS+ was still evident for control ants and ants treated with 20mM epinastine (wilcoxon tests p < 0.001), and still absent in ants treated with 100mM epinastine (wilcoxon test p = 0.77). There was only a trend for flupentixol-treated ants to discriminate CS+ and CS0 (p = 0.092; Figure 5B). However, all ants appeared to be affected by the receptor blockers, as the preference indices of all ants treated with receptor blockers were lower than those of the control ants (glmm, all p < 0.05, Tab. S3).

To test whether ant learning or memory retrieval may be prevented by a general lethargy caused by the receptor blockers, we tested whether they affected the walking speed of the ants. Indeed, during the first retention test, ants treated with both concentrations of epinastine were slower than control ants (Fig. 6A; glmm p < 0.01; Tab. S4), but flupentixol did not affect walking speed (p = 0.29). However, the speed reduction was only in the range of 20% and should not have prevented the ants from encountering the odours. Indeed, only a single ant, treated with 100mM epinastine, did not enter the quadrant with the rewarded odour during the retention test. One day later, only the walking speed of epinastine 100mM ants was reduced (glmm epinastine p < 0.05; both other treatments p > 0.59; Tab. S4; Fig. 6B).

**Figure 6.**
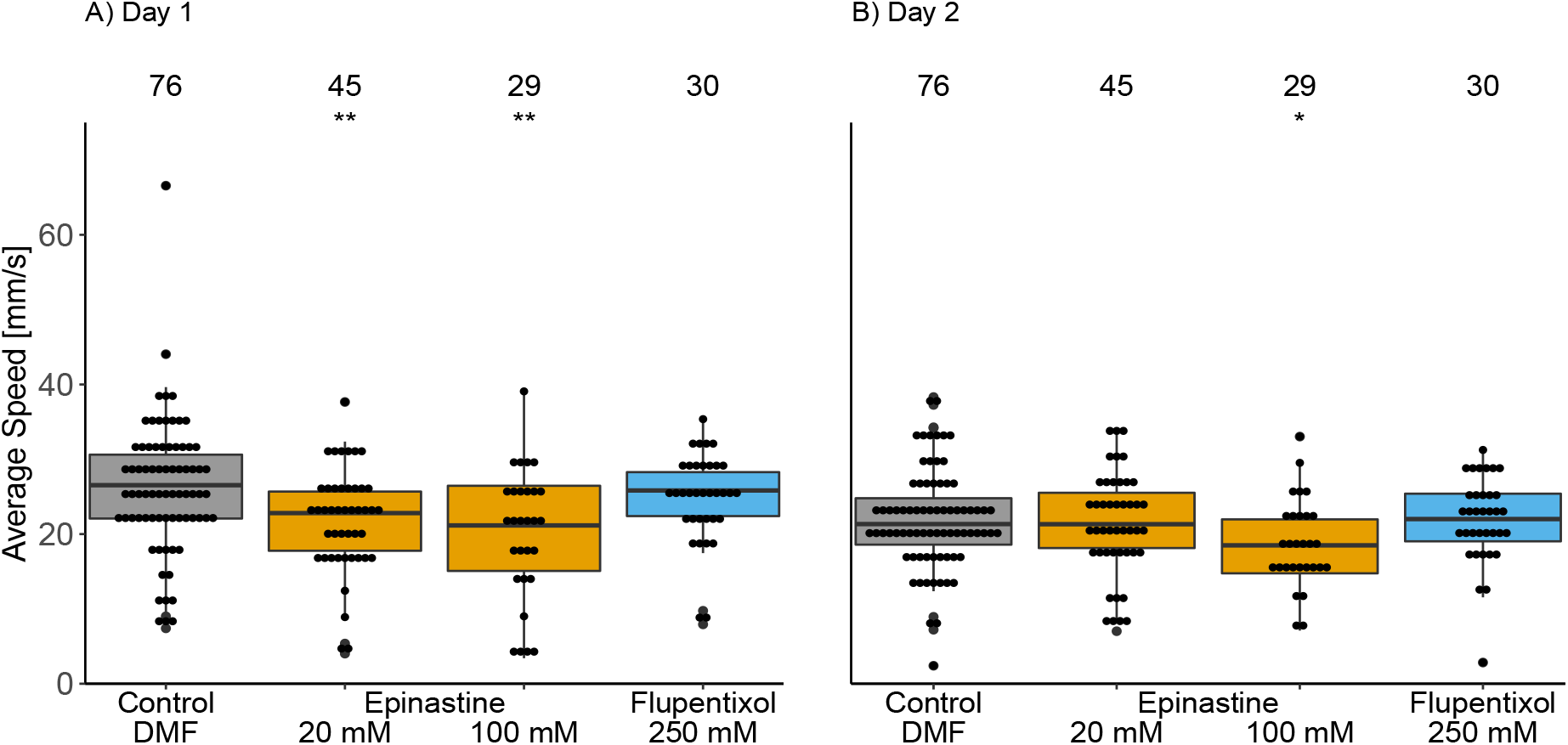
Walking speed during retention tests of control ants and ants that topically received either epinastine or flupentixol. Stars indicate treatments that changed the average speed significantly compared to the control treatment (glmm * p < 0.05; ** p < 0.01).

## Discussion

Using a classical conditioning protocol, we could show that black garden ant workers can learn to associate blends of linear hydrocarbons with a sucrose reward, and that the memory is stable for at least one day. Exposing the ants to the octopamine receptor blocker epinastine prevented memory formation or retrieval. In contrast, when we exposed the ants to the dopamine receptor blocker flupentixol, the short term memory persisted but the long-term memory was affected. Both octopamine and dopamine signalling are thus involved in the formation of appetitive memory in ants. Our results further suggest that the active period of the receptor blockers is limited in that learning is possible again a few hours after blocker ingestion.

### Associating hydrocarbons with a sucrose reward

During the test trials, the ants spent more time on the quadrant that contained the conditioned stimulus (CS+) than that containing a neutral stimulus not previously associated with a sucrose reward (CS0). The increase in time spent in the CS+ quadrant is likely a result of the ants searching for the reward. The ants are thus able to learn and discriminate blends of linear hydrocarbons. Most appetitive conditioning assays in hymenoptera rely on volatile odours that are typically emitted from flowers, to closely resemble the natural situation in which learning can lead to flower constancy in bees (Koethe et al., 2020). However, many pheromones of social insects are less volatile hydrocarbons. Our study confirms previous work on Carpenter Ants and Argentine Ants that demonstrates hydrocarbons are also adequate stimuli for appetitive conditioning, and that the meaning of pheromones and signature mixtures could potentially be changed through learning (Bos et al., 2012; van Wilgenburg et al., 2012; di Mauro et al., 2015; Sharma et al., 2015).

### Involvement of octopamine in appetitive learning

Unlike control treated ants, ants that were treated with 100mM epinastine topically to the thorax did not prefer the conditioned hydrocarbon odour over a non-conditioned odour within five minutes after the learning trials, and neither did they prefer the conditioned stimulus after a break of one day. As epinastine blocks octopamine receptors, this shows that octopamine signalling is required for the formation of short and long term memory of appetitive stimuli in ants, potentially because octopaminergic neurons relay the unconditioned stimulus to the Kenyon cells (e.g. VUMx1 in bees, Rein et al., 2013). Our findings confirm previous work demonstrating such an effect in fruit flies, honey bees, and crickets (Schwaerzel et al., 2003; Giurfa, 2007; Mizunami and Matsumoto, 2017).

Epinastine does not only affect memory but potentially also other traits. Under the influence of epinastine, silkmoths were less sensitive for pheromones (Pophof 2000), the metabolism of hungry honeybees was reduced (Buckenmüller et al.,2017), and aggression was modulated in crickets and red wood ants (Rillich and Stevenson 2011, 2018; Yakovlev, 2018). In our study, the mobility of ants treated with epinastine was reduced during the first retention test. For the high dose of epinastine, this effect was still visible one day later, which may indicate that epinastine is only slowly degraded. However, the ants still visited all quadrants of the arena during the retention tests, and they also reacted to the sucrose solution during the learning trials. We thus argue that the failure to identify the CS+ in the retention tests is not due to a decrease of motivation or even an inability to walk, but due to a lack of memory formation or retrieval. In fact, in all our learning trials with epinastine-treated ants, the time until the ants found the sucrose solution decreased. A simple explanation would be that the ants do learn to expect sucrose in the arena and search for it, which decreases search time, but they do not associate it with the odour and thus do not spend more time with the previously rewarded odour in the non-rewarded retention trials.

Ants topically treated with a lower dose of epinastine were not impaired in their learning ability. More interestingly, the effects were still visible when we fed epinastine with the food, but weaker than when we directly applied the drug to the thorax. It is possible that only part of the dose is entering the hemolymph when the drugs are taken up with the food. Ant workers possess a crop used to store food that is later regurgitated to feed nestmates (Greenwald et al., 2018). Some epinastine entering the crop may thus never reach the octopamine receptors in the brain, exposing them to lower doses than intended. It may take a while until epinastine is taken up into the hemolymph and reaches the brain, meaning that it will take action only after a certain delay (but see Barron et al., 2007: octopamine delivered through the food reaches the honey bee brain in less than one hour). We thus fed the ants epinastine at different intervals before subjecting them to the learning trials. However, longer intervals only reduced the effect of epinastine, and ca. 20 hours after being fed the receptor blockers, the preference indices of epinastine-fed individuals were no different than those of control ants. Finally, when ants were fed epinastine, we found only weak effects on the short term memory retention. Given that there was a clear effect on the short term memory when epinastine was topically applied, this might have to do with long and short term memory being formed in different parts of the brain (e.g. different populations of Kenyon cells, Sachse and Galizia, 2003) and potentially at different times. If short term memory formation is quicker than long term memory formation, epinastine may have reached the brain before the long term memory formation, but after short term memory formation was complete.

### Involvement of dopamine

In our experiments, flupentixol strongly affected the long term memory as well, but not necessarily the short term memory. After topical application, we did not find any evidence of flupentixol affecting the short term memory. Flupentixol blocks dopamine receptors, so that our observation suggest that the dopaminergic neurons are probably not required in ants to relay the unconditioned stimulus to the Kenyon cells (as it is in fruit flies and bees, Schwaerzel et al., 2003; Giurfa, 2007). However, we found weak effects of flupentixol on the short term memory when we fed it to the ants, which may indicate that a functional dopaminergic system may improve learning performance.

In contrast to the short term memory, the long term memory was always affected when we administered flupentixol via either food or topical application. It is particularly curious that this even happened after topical application of flupentixol, when ants still clearly preferred the rewarded stimulus in the short term memory retention tests. This indicates a necessary involvement of dopaminergic neurons in the consolidation phase of long term memory formation that is independent of the short term memory and cannot be rescued by octopaminergic neurons, which are still functioning when only flupentixol is administered. The dopaminergic neurons may thus not be relaying the unconditioned stimulus to the learning centres, but be part of feedback loops or modulation pathways (Eschbach et al., 2020). Long term memory formation is a more complex process than short term memory formation (e.g. Menzel, 1999; Villar, 2020) and memory consolidation might rely on additional neural circuits that could be dopaminergic. For example, some lateral neurons in the antennal lobes are dopaminergic (Galizia and Sachse, 2010) and might be used to improve the perception of relevant odours such as the CS+. In the mushroom body, Kenyon cells receive feedback from the primary protocerebral anterior cluster via dopaminergic neurons in fruit fly larvae (Lyutova et al., 2019).

Blocking dopamine receptors does not only influence appetitive and aversive learning. Blocking the AmDOP2 receptor has led to reduced motor behaviours and an increase of grooming in honey bees (Mustard et al., 2010). In *Drosophila*, social interactions and temperature preferences seem to be affected (Verlinden, 2018). In our study, there was no evidence of effects on motor behaviour, since walking speed of flupentixol-treated ants did not differ from that of control ants. Interestingly though, flupentixol fed 5-8 hours before learning, the treatment that affected both the short and the long term memory, also prevented an improvement of the times the ants needed to find the reward during the learning trials (Fig. 2). Also, in the flupentixol-treated groups, particularly many individuals did not find the sucrose reward during the two minutes of the learning trial (Tab. S1). One possible explanation could be that blocking dopamine signalling interferes with the motivational state of the ants. If lack of dopamine signalling would make them believe they were not hungry, they might not be primed to seek the sucrose reward in the learning trials, but may still be drawn to the conditioned stimulus in the retention tests because of its familiarity. Indeed, dopamine is important for hunger signalling in *Drosophila* (Krashes et al. 2009; Siju et al., 2021). Hunger affects the motivational state, which is important for memory formation and retrieval because satiated flies do not seek food. However, dopaminergic neurons appear to signal satiation rather than hunger in *Drosophila*, so that blocking them should improve motivation and thus memory formation (Krashes et al., 2009). If the neural circuits were similar in ants, hunger could thus not explain the curious pattern caused by blocking dopamine receptors. However, the motivation of ants as social beings is perhaps regulated less by their personal physiological hunger and more by that of their colony, so that entirely different regulatory circuits might be involved here (e.g. Cholé et al., 2019). In any case, such a lack of hunger or motivation could only explain an effect of flupentixol on the short term memory in the feeding trial: In the topical application experiment, ants did initially learn in learning trials and remembered in the first retention test, but clearly failed to remember on the next day. This observation indicates an involvement of dopamine in the consolidation of the long term memory, independent short term memory formation and motivation.

## Conclusions

*Lasius niger* ants can learn to associate mixtures of linear hydrocarbons to a sugar reward and remember the association for at least 24 hours. Both octopamine and dopamine appear to be involved in memory formation, with strong effects of a pharmacological octopamine knockout on both the short and the long term memory. Dopamine signalling is required for the formation of the long term memory but not necessarily for the short term memory.

## Acknowledgements

We would like to thank the members of the Department of Ecology and Evolution, University of Freiburg, Germany, and Andrew Straw for fruitful discussions. Thanks to Mélanie Bey for helpful comments on a previous version of the manuscript, and to the German Research Foundation (DFG grant) for funding.

