## Supplementary Information for "Appetitive learning relies on octopamine and dopamine in ants"

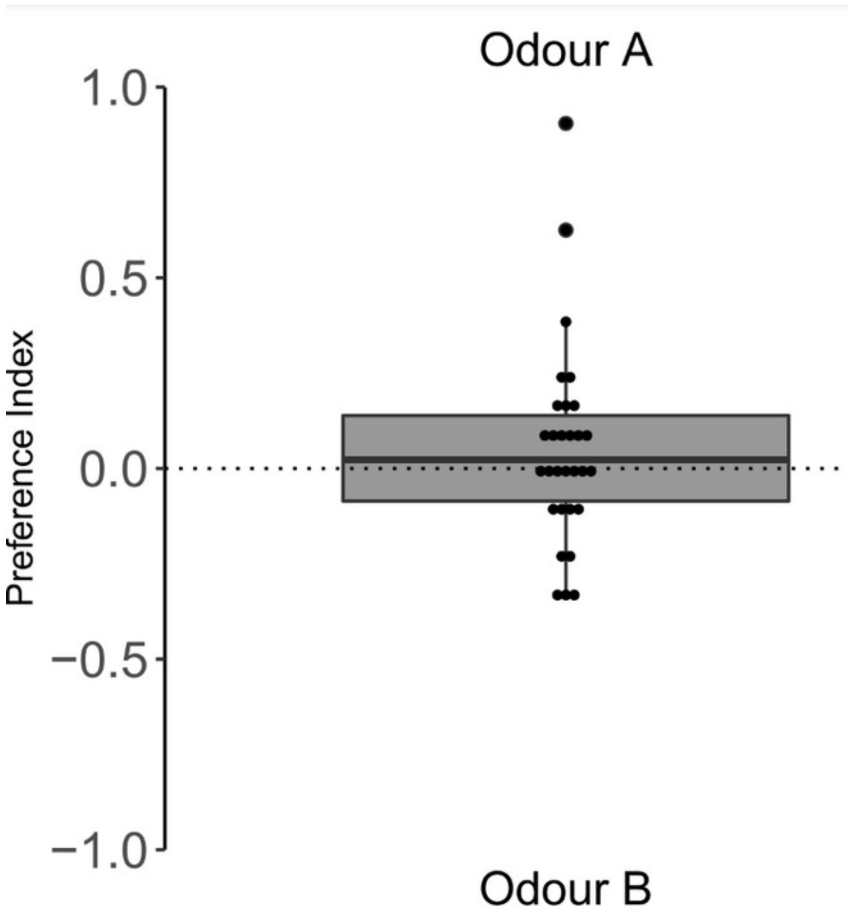

**Supplementary Figure 1**: The preference index of untrained ants for odour mixture A over odour mixture B when presented on glass slides. The odour mixtures were later used as rewarded stimulus (CS+) and unrewarded stimulus (CS0) in the learning experiments. Both odour mixtures contained equal proportions of three n-alkanes (odour A: n-C18, n-C21, and n-C27; odour B: n-C20, n-C22, n-C25), and the ants did not prefer either (n = 30, wilcoxon test V = 253, p = 0.69).

**Supplementary table 1:** The number of ants that entered the learning trials, and the sample size of the following retention tests after removing ants that did not find the sugar solution during at least two learning trials, or died before the second retention test.

| Experiment | Treatment | Sample Size | Sample Size – Adjusted |
|---|---|---|---|
| Application | DMF – Control | 76 | 76 |
| | Epinastine 20mM | 45 | 45 |
| | Epinastine 100mM | 29 | 29 |
| | Flupentixol 250mM | 32 | 30 |
| Feeding | Untreated – Control | 30 | 30 |
| | Epinastine 1-3 Hours | 30 | 27 |
| | Epinastine 5-8 Hours | 29 | 29 |
| | Epinastine 17-26 Hours | 27 | 25 |
| | Flupentixol 1-3 Hours | 28 | 26 |
| | Flupentixol 5-8 Hours | 25 | 21 |

**Supplementary table 2:** glmm on preference indices during retention tests of ants fed with receptor blockers.

| Response Variable | Predictor | Post. Mean | l-95% CI | U-95% CI | Effect Sample | pMCMC |
|---|---|---|---|---|---|---|
| Preference Index Day 1 | Epinastine 1-3 Hours | -0.02408 | -0.13602 | 0.09734 | 4100 | 0.688 |
| | Epinastine 5-8 Hours | -0.10452 | -0.21775 | 0.00805 | 4925 | 0.073 |
| | Epinastine 17-26 Hours | -0.07946 | -0.19765 | 0.03705 | 4998 | 0.186 |
| | Flupentixol 1-3 Hours | -0.11920 | -0.23403 | 0.00010 | 4700 | 0.044 |
| | Flupentixol 5-8 Hours | -0.08786 | -0.21319 | 0.03606 | 4700 | 0.160 |
| Preference Index Day 2 | Epinastine 1-3 Hours | -0.09872 | -0.19858 | 0.18057 | 4491 | 0.063 |
| | Epinastine 5-8 Hours | -0.02446 | -0.12837 | 0.07362 | 4700 | 0.635 |
| | Epinastine 17-26 Hours | 0.02232 | -0.08314 | 0.12693 | 4700 | 0.671 |
| | Flupentixol 1-3 Hours | -0.03283 | -0.13307 | 0.07369 | 4700 | 0.537 |
| | Flupentixol 5-8 Hours | -0.03771 | -0.14776 | 0.07340 | 4359 | 0.509 |

**Supplementary table 3:** glmm on preference indices during retention tests of ants that received a topical application of receptor blockers.

| Response Variable | Treatment | Post. Mean | l-95% CI | U-95% CI | Effect Sample | pMCMC |
|---|---|---|---|---|---|---|
| Preference Index Day 1 | Epinastine 20mM | -0.04108 | -0.1600 | 0.06608 | 4960 | 0.447 |
| | Epinastine 100mM | -0.22515 | -0.3563 | -0.09125 | 4551 | < 0.001 |
| | Flupenthixol 250mM | -0.09118 | -0.2253 | 0.03194 | 4700 | 0.173 |
| Preference Index Day 2 | Epinastine 20mM | -0.10816 | -0.2108 | -0.00704 | 4903 | 0.043 |
| | Epinastine 100mM | -0.41991 | -0.5392 | -0.30225 | 4700 | < 0.001 |
| | Flupenthixol 250mM | -0.23913 | -0.3584 | -0.11913 | 4496 | < 0.001 |

**Supplementary table 4**: glmm on the walking speed of ants in the retention tests after topical application of receptor blockers.

| Response Variable | Treatment | Post. Mean | l-95% CI | U-95% CI | Effect Sample | pMCMC |
|---|---|---|---|---|---|---|
| Average Speed Day 1 | Epinastine 20mM | -4.35 | -7.215 | -1.446 | 4501 | 0.002 |
| | Epinastine 100mM | -5.97 | -9.362 | -2.341 | 2361 | 0.001 |
| | Flupentixol 250mM | -1.81 | -5.046 | 1.594 | 4700 | 0.294 |
| Average Speed Day 2 | Epinastine 20mM | -0.597 | -2.790 | 1.616 | 4700 | 0.597 |
| | Epinastine 100mM | -2.952 | -5.574 | -0.289 | 6173 | 0.031 |
| | Flupentixol 250mM | -0.512 | -3.172 | 2.124 | 4700 | 0.713 |
